# Population structure and genetic diversity of smooth newts (*Lissotriton vulgaris*) in North Tyrol, Austria: influences of allochthonous individuals and conservation implications

**DOI:** 10.64898/2026.01.23.701301

**Authors:** Katharina Theresa Stonig, Marlene Haider, Florian Glaser, Florian M. Steiner, Birgit C. Schlick-Steiner

## Abstract

Amphibians are threatened worldwide by various environmental and anthropogenic factors, making non-invasive conservation studies particularly valuable. Newts are one example of a thus challenged amphibian group. In Austria, local population declines of newts have been observed, with the smooth newt (*Lissotriton vulgaris*) being strongly affected. In this study, skin swabs were used as a non-invasive method to gather DNA, combined with established microsatellite markers. We sampled 139 *L. vulgaris* individuals at ten sites in North Tyrol, Austria, and, for comparison, 22 *L. vulgaris meridionalis* individuals in Brixen, Italy. We genotyped all individuals and analysed their population structure. We demonstrate the presence of three distinct *L. vulgaris* population clusters and find differences in population structure between supposedly introduced allochthonous *L. vulgaris* individuals and autochthonous populations, as evidenced by differences in Bayesian clustering and elevated values of the fixation index F_ST_. A captive population in a zoological garden, with origins in the Kramsacher Loar in the Tyrolean Unterland (eastern part of Tyrol), performed poorly in terms of conservation genetics, with low genetic diversity (number of alleles) and clear genetic differentiation from populations in the wild (high pairwise F_ST_ values with wild individuals, clear separation in cluster analysis). Habitat restoration programs are a crucial aspect of amphibian conservation, as they restore ecosystems that are critical to the animals’ survival. While breeding programs can play an additional role in the future, they must carefully consider genetic diversity to ensure resilient and viable populations, especially in the face of climate change and chytrid fungus infection. This study emphasizes the significance of considering the geographic origin and genetic diversity of newts in conservation efforts. It also serves as a foundation for future population genetic studies of newts in Austria.

## Introduction

Salamandridae, also known as tailed amphibians, are a distinct family within the class Amphibia. One group within the Salamandridae are newts, characterized by their multiphasic lifecycle, the occurrence of neoteny, and limited dispersal ability (Malmgren 2001; Sparreboom 2014). Of the six native newts in Austria (Glandt 2018), four species are documented in Tyrol: the crested newt *Triturus cristatus*, the Italian crested newt *Triturus carnifex*, the alpine newt (*Mesotriton alpestris*), and the smooth newt (*Lissotriton vulgaris*) (Kofler 1978; Kostenzer et al. 1996; Glaser et al. 2006; Glaser 2008) The alpine and the smooth newt are the most commonly found newt species in Tirol and are widespread throughout Austria. Both newt species occur syntopically at lower elevations and share their habitat in ponds and pools without fish stocks. Temporary pools, such as ruts in forest tracks, are also often used as spawning sites (Sparreboom 2014; Kopecký 2023). In all life stages, newts are strictly carnivorous, feeding on zooplankton, diverse insects and their larvae, invertebrates, and tadpoles, and are cannibalistic (Sparreboom 2014). Lower elevations in the valleys are preferred by *L. vulgaris*, the less common newt species found in Tyrol

Like all amphibians, newts are highly vulnerable to environmental impacts, such as habitat loss and fragmentation, agricultural destruction, climate change as well as the spreading of the chytrid fungi (Jehle and Arntzen 2002; Sparreboom 2014; Buono et al. 2018; Longo et al. 2019). *Lissotriton vulgaris* is classified as LC (of least concern) in the IUCN Red List assessment (Species Survival Commission Species Survival Commission 2021; Species Survival Commission 2022) but suffers from regional extinctions throughout Europe and is classified as NT (near threatened) at the national level in Austria (Gollmann 2007). The Tyrolean Inn Valley in the Austrian Alps (Figure 1) poses a particular challenge with its limited space and extensive human land use (Zimmermann et al. 2010), which fragments and limits amphibian habitats. A fragile population trend has been evident since the early 1990s in *L. vulgaris* populations, marked by pronounced fluctuations in the number of individuals across various sites, including regions with autochthonous populations such as that in the Kramsacher Loar (Figure 1), in the east of the Tyrolean Inn Valley (Landmann et al. 1999; Landmann and Fischler 2000; Glaser et al. 2003; Glaser 2008). This is most likely caused due to fragmentation by roads and infrastructure, which has a particularly massive impact in a linear, densely populated Alpine valley, as well as to fish stocking and agricultural pressure (Glaser 2008; Glandt 2018). Countermeasures have been taken in recent decades to combat the few remaining autochthonous populations, but these have been characterized by questionable decisions, such as the restocking of newts with allochthonous individuals from a narrow gene pool. Populations resulting from releases and involuntary introductions have been documented for this region (Glaser et al. 2006). In the case of amphibians in particular, there is a need for targeted, science-based conservation measures (Nori et al. 2015).

**Figure 1.**
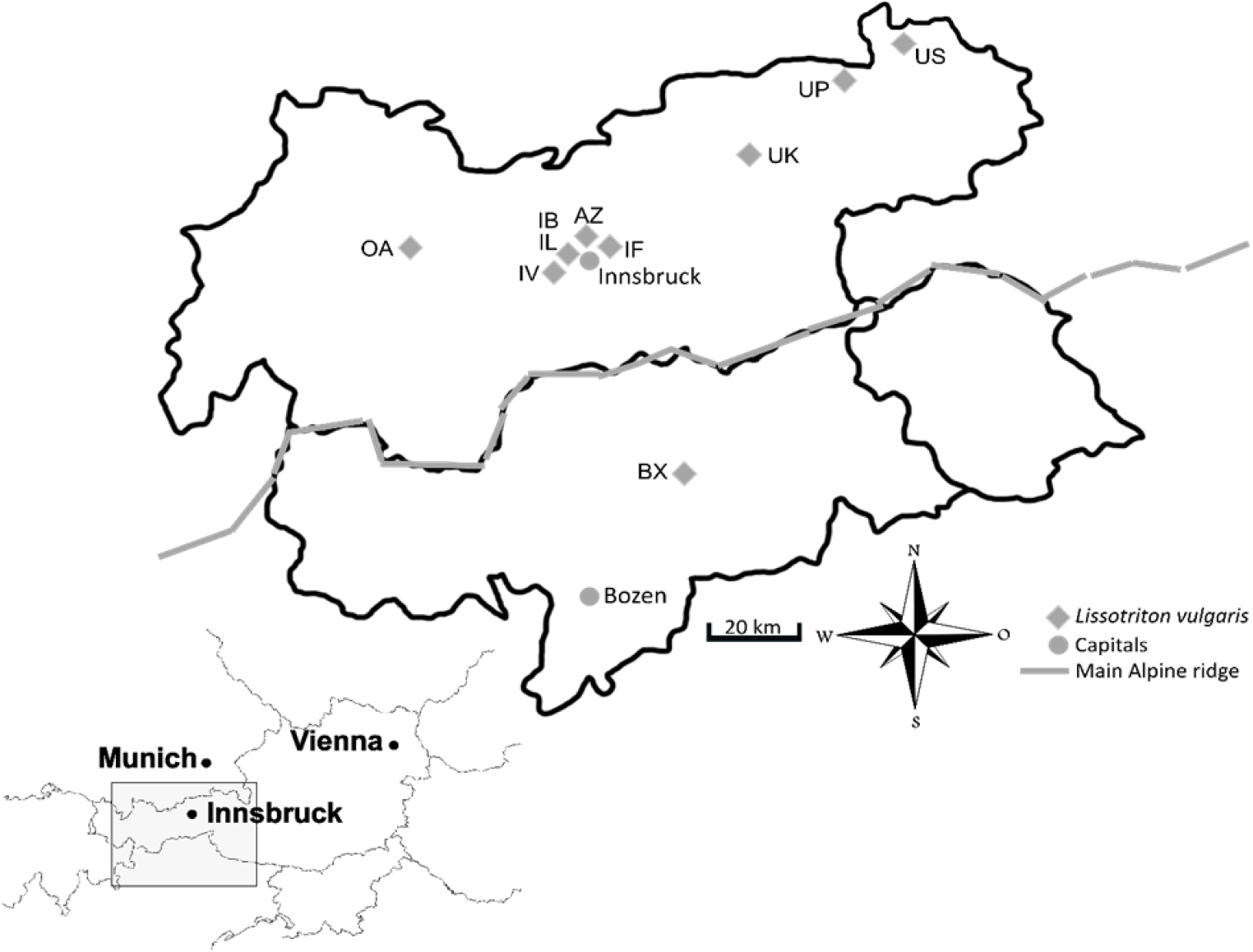
Schematic illustration of the study area in North and South Tyrol; pentagons represent sampling sites of *Lissotriton vulgaris*; dotted lines suggest the main Alpine ridge as a special barrier; Abbreviations of sampling sites: Oberland Area47 (OA), Innsbruck Völser Teich (IV), Innsbruck Bauteil 8 (IB), Innsbruck Lohbach (IL), Innsbruck Fuchsloch (IF), Alpenzoo Innsbruck (AZ), Unterland Kramsacher Loar (UK), Unterland Pfrillsee (UP), Unterland Schwemm (US), Brixen Millander Au (BX).

*Lissotriton vulgaris* has previously been the focus of population genetic studies, and microsatellite markers have already been developed (Buono et al. 2018). Microsatellite markers are a popular tool for analysing population structure, population ecology, and conservation status (Ashley and Dow 1994; Jehle and Arntzen 2002; Buono et al. 2018). Combined with non-invasive skin swabs, this approach has great potential for minimally invasive studies of sensitive, protected amphibian species – as many amphibians are strictly protected, research must not cause additional harm to individuals and populations (Prunier et al. 2012). This technique of using microsatellite analysis and skin swabs has been tested on frogs and salamanders by research groups with different degrees of success (Prunier et al. 2012; Ringler 2018), concluding that dry amphibian skin (particularly when sampled ventrally) negatively affects the quality of the DNA obtained. More deliberate rubbing of grainier skin produced satisfactory results (Prunier et al. 2012).

Here, we aim to contribute further insights by applying non-invasive methods to analyse newts native to Tyrol, Austria, given the mixed success of previous research on amphibians, particularly with regard to the quality of DNA obtained (Ringler 2018). We analysed 161 *L. vulgaris* individuals. We compared autochthonous populations with sites where allochthonous individuals (with origins from a location in East-Tyrol, approximately 80 km in a straight line from the closest autochthonous population) had been artificially introduced. We sampled genetically isolated individuals (originating from the Kramsacher Loar, in the Tyrolean Unterland) from the zoological garden Alpenzoo Innsbruck and compared them with individuals from their origin site. As a population-genetic outgroup, we also sampled *Lissotriton vulgaris meridionalis* in Brixen, Italy, a *Lissotriton* subspecies with a distinct evolutionary lineage (Maura et al. 2014). We calculated common population genetic indices for *L. vulgaris* and analysed the population structure using Bayesian clustering. With this study, we gain a better understanding of the current population structures of *L. vulgaris* in the Tyrolean Inn Valley, shed light on population genetic isolation processes, and learn to better understand the consequences of the introduction of allochthonous individuals.

## Materials and Methods

### Sampling

Sampling of *L. vulgaris* was conducted at night time between April and September 2022 at ten locations in Tyrol, Austria (Oberland Area47 (OA), Innsbruck Völser Teich (IV), Innsbruck Bauteil 8 (IB), Innsbruck Lohbach (IL), Innsbruck Fuchsloch (IF), Alpenzoo Innsbruck (AZ), Unterland Kramsacher Loar (UK), Unterland Pfrillsee (UP), Unterland Schwemm (US)) and at one location in South Tyrol, Italy (Brixen Millander Au (BX)) (Figure 1).

A total of 161 newts were collected using landing nets of different sizes and buckets. Species identification according to Thiesmeier and Franzen (2018) and sexing of the individuals were conducted morphologically on site. To prevent cross-contamination of DNA and potential diseases, each individual was handled with new gloves. All equipment was disinfected after each use with commercially available chlorine-based bleaching agent (concentration <5%). Non-invasive skin swabs (HEINZ HERENZ Medizinbedarf GmbH dry swabs) were used for sampling. The ventral and both lateral sides as well as the cloacal region toward the hindlegs were swiped ten times per individual to obtain a sufficient amount of tissue for DNA extraction. Each individual was handled as briefly as necessary. The skin swabs were stored immediately after sampling in a freezer at −20 °C and transferred to −70 °C the next day.

### Analysis

The QIAamp® DNA Micro Kit (QIAGEN GmbH, Hilden, Germany) was used for DNA extraction, following the manufacturer’s protocol with specific modifications introduced between steps 4 and 5: Swabs were centrifuged for 1 min at 14,000 rpm through a DNA IQTM Spin Basket into 1.5 ml standard Eppendorf tubes. This step was repeated twice, with a final centrifugation for 2 min at 14,000 rpm. The baskets were removed, and the tips were incubated at 56 °C for 20 min. DNA was eluted in 40 μl distilled water. The DNA extracts were stored at −20 °C.

Eleven microsatellite markers were used for *L. vulgaris* (Th09, Tv12, Tv3Ca9, Lm_521, Lm_528, Lm_749, Lm_870, LVG-210, LVG-542, Lm_013, Tv3Ca19) (Buono et al. 2018). PCRs were performed with a final reaction volume of 5 μl. For validation of PCR results, the multiplex PCRs were performed in triplicate. To enhance the PCR output, 5% DMSO und BSA (20 ug/μl) were added to the GoTaq® Green Master Mix (Promega Corporation, Madison, USA).

Five different cycling conditions were employed in this study. All programs included an initial denaturation period of 15 min at 95 °C, an elongation period of 45 s at 72 °C, and a final extension period of 10 min at 72 °C. **Program 1:** 37 cycles of denaturation at 95 °C for 45 s, annealing at 61 °C for 45 s, used for Lm_521, Lm_528 **Program 2:** 37 cycles of denaturation at 95 °C for 45 s, annealing at 55 °C for 45 s, used for Th09, Tv3Ca9, Tv3Ca19, Lm_013, LVG-210 **Program 3:** 37 cycles of denaturation at 95 °C for 45 s, annealing at 57 °C for 45 s, used for Tv12 **Program 4:** 40 cycles of denaturation at 95 °C for 45 s, annealing at 57 °C for 45 s, used for Lm_749, Lm_870 **Program 5:** 40 cycles of denaturation at 95 °C for 45 s, annealing at 56 °C for 45 s, used for LVG-542. Each PCR was repeated three times for each sample to ensure correct genotyping (Taberlet et al. 1996). The PCR products were then sent to the DNA Sequencing & Genotyping Facility at the University of Chicago Comprehensive Cancer Center (Chicago, USA) and Ingenetix GmbH (Haidingergasse 1, 1030 Vienna, Austria) for electrophoretic sizing via capillary electrophoresis. GeneMarker v2.7.0/3.0.1 (SoftGenetics, State College, PA, USA) was used for subsequent allele calling.

After removal of markers which had failed (mismatching alleles in repeated runs or monomorphism), the following eight microsatellite markers were used for population genetic analysis: Th09, Tv12, Tv3Ca9, Lm_749, Lm_870, LVG-542, Lm_013, Tv3Ca19 (Supplementary Table S1). To prevent the results from being distorted by high numbers of missing values, individuals with a minimum of four called loci were selected, leaving 121 individuals. Both the complete (161 individuals) and the selected (121 individuals) dataset were analysed.

All calculations were performed using R v4.2.2 (R Core Team 2022). Linkage disequilibrium (LD) was assessed using poppr v2.9.5 (Kamvar et al. 2015). Null alleles were tested with the PopGenReport v3.1. (Gruber and Adamack 2015), and observed and expected heterozygosity (Ho and He) were calculated using adegenet v2.1.10 (Jombart 2008). Bartlett’s K^2^ test was used to compare the variance between Ho and He, integrated in the package stats v4.2.2. The pairwise fixation index (F_ST_) and the inbreeding coefficient (F_IS_) were calculated using hierfstat v0.5-11 (Goudet et al. 2022). Using STRUCTURE v2.3.4 (Jul 2012), population structure was inferred using a Bayesian cluster analysis (Pritchard et al. 2000) with an admixture approach with 1,000,000 generations burn-in and 2,000,000 generations sampling phase and 10 replicates. The most likely numbers of populations were identified using ΔK (Evanno et al. 2005) using pophelper v2.3.1 (Francis 2017).

## Results

A total of 161 *Lissotriton vulgaris* samples (87 females, 54 males, and 20 juveniles) were analysed, originating from all ten locations sampled (Figure 1), with an average of 16 individuals per site (min = 1, max = 23; Table 1).

**Table 1.** Sampling sites (site), number of sampled *Lissotriton vulgaris* individuals (N), allele number (Na), mean expected heterozygosity (He), mean observed heterozygosity (Ho), 95% confidence interval (Ho Median (95%CI)), number of microsatellite marker used for calculation of He and Ho (n) and Bartlett’s K square test of variance, as a proxy of differences between He and Ho (Bartlett’s K^2^) with p values in brackets. Calculations for UK were not possible. AZ results for He and Ho should not be considered, since Bartlett’s K^2^ is significant. Abbreviations for different sites can be taken from Figure 1.

| Site | N | Na | He (± SD) | Ho (± SD) | Ho Median (95% CI) | n | Bartlett's $K^2$ |
| --- | --- | --- | --- | --- | --- | --- | --- |
| OA | 23 | 39 | 0.59 (±0.25) | 0.35 (±0.34) | 0.23 (0.11-0.56) | 8 | 0.65 (0.41) |
| IV | 10 | 27 | 0.58 (±0.17) | 0.46 (±0.29) | 0.44 (0.33-0.57) | 8 | 1.79 (0.18) |
| IB | 22 | 27 | 0.61 (±0.27) | 0.21 (±0.23) | 0.09 (0.04-0.34) | 7 | 0.13 (0.72) |
| IL | 22 | 24 | 0.62 (±0.26) | 0.23 (±0.25) | 0.10 (0.05-0.41) | 7 | 0.00 (0.99) |
| IF | 13 | 28 | 0.52 (±0.23) | 0.32 (±0.23) | 0.29 (0.21-0.40) | 8 | 0.00 (0.97) |
| AZ | 23 | 24 | 0.55 (±0.14) | 0.52 (±0.34) | 0.43 (0.29-0.71) | 8 | 4.82 (0.03) |
| UK | 1 | 1 |  |  |  |  |  |
| UP | 5 | 19 | 0.62 (±0.37) | 0.46 (±0.46) | 0.38 (0.06-0.88) | 6 | 0.28 (0.59) |
| US | 20 | 37 | 0.58 (±0.38) | 0.30 (±0.25) | 0.30 (0.14-0.42) | 7 | 1.12 (0.29) |
| BX | 22 | 55 | 0.67 (±0.24) | 0.48 (±0.41) | 0.48 (0.15-0.81) | 8 | 1.69 (0.19) |

We did not find any linkage between or null alleles in the eight loci tested. The Innsbruck sites (AZ, IL, and IB) had the lowest numbers of alleles (Na= 24,24,27) compared with the number of samples (N=23,22,22). The number of markers used to calculate the expected and observed heterozygosity differed across the ten sampling sites. AZ was the only site where the number of makers used decreased, from eight to six, for the selected dataset (Table 1, Supplementary Table S2). Median observed heterozygosity (Ho Median (95%CI)) ranged from 0.09 at IB to 0.48 at BX (Table 1). The sites IL and IB, located in Innsbruck, had the lowest values of Ho in both the complete and the selected dataset (Table S2).

The mean squared distance (d^2^) was generally low and similar for most sites, except for BX, which showed the highest values (Figure 2). The lowest d^2^ values were observed in both sites in the Tyrolean Unterland (UP, US). A similar pattern was observed in the selected dataset, with one exception at AZ, where the values were lower (which was already seen in the Ho values) (Supplementary Figure S1). The patterns found were mostly in line with the multi-locus heterozygosity (MLH), which was highest for BX, and IV also showed higher values (Figure 2, Figure S1). F_IS_ values showed high variance with a tendency toward positive values across all sampling sites (Figure 2, Figure S1). Looking at the selected dataset, the variance was even higher; however, all sites but UP had a positive lower 95% confidence interval limit, again demonstrating tendencies for overall positive values. Pairwise F_ST_ values ranged from 0.04 between IL and IF to 0.32 between AZ and US. The highest F_ST_ values were observed between AZ and the other sampling sites. The comparison of the complete *L. vulgaris* dataset (161 individuals; Table 2) with the selected dataset (121 individuals; Supplementary Table S3) revealed no major difference.

**Table 2.** Pairwise fixation index (F_ST_) values between ten different sampling sites of *Lissotriton vulgaris*. Values that exceed 0.05 suggest low genetic connectivity between the sites. Small numbers below the F_ST_ values represent the lower and upper 95% confidence interval. Values highlighted in bold are significant. AZ shows the highest genetic isolation with all sampling sites. Abbreviations for different sites can be taken from Figure 1.

|  | OA | AZ | IB | BX | IF | UK | IL | UP | US |
| --- | --- | --- | --- | --- | --- | --- | --- | --- | --- |
| AZ | **0.31**<br>0.16; 0.47 |  |  |  |  |  |  |  |  |
| IB | **0.15**<br>0.03; 0.33 | **0.21**<br>0.08; 0.40 |  |  |  |  |  |  |  |
| BX | **0.12**<br>0.06; 0.18 | **0.27**<br>0.12; 0.42 | **0.19**<br>0.11; 0.33 |  |  |  |  |  |  |
| IF | **0.10**<br>0.01; 0.19 | **0.25**<br>0.02; 0.48 | 0.08<br>-0.03; 0.20 | **0.18**<br>0.12; 0.24 |  |  |  |  |  |
| UK | 0.06<br>-0.04; 0.17 | **0.10**<br>0.01; 0.22 | 0.04<br>-0.00; 0.12 | 0.04<br>-0.09; 0.14 | **0.10**<br>0.02; 0.22 |  |  |  |  |
| IL | **0.10**<br>0.03; 0.14 | **0.23**<br>0.10; 0.41 | 0.05<br>-0.01; 0.12 | **0.15**<br>0.09; 0.19 | **0.04**<br>0.00; 0.08 | 0.07<br>-0.02; 0.17 |  |  |  |
| UP | 0.01<br>-0.10; 0.06 | 0.21<br>-0.01; 0.46 | 0.03<br>-0.02; 0.08 | 0.09<br>-0.01; 0.21 | 0.01<br>-0.14; 0.15 | 0.01<br>-0.20; 0.25 | **0.07**<br>0.01; 0.10 |  |  |
| US | 0.02<br>-0.05; 0.07 | **0.32**<br>0.09; 0.56 | **0.17**<br>0.01; 0.41 | **0.13**<br>0.05; 0.22 | 0.06<br>-0.03; 0.14 | 0.07<br>-0.07; 0.25 | **0.07**<br>0.03; 0.13 | 0.04<br>-0.15; 0.04 |  |
| IV | **0.19**<br>0.04; 0.37 | **0.20**<br>0.03; 0.36 | **0.12**<br>0.04; 0.21 | 0.17<br>-0.00; 0.35 | **0.18**<br>0.08; 0.28 | 0.02<br>-0.17; 0.23 | **0.12**<br>0.06; 0.19 | 0.04<br>-0.05; 0.14 | **0.20**<br>0.04; 0.43 |

**Figure 2.**
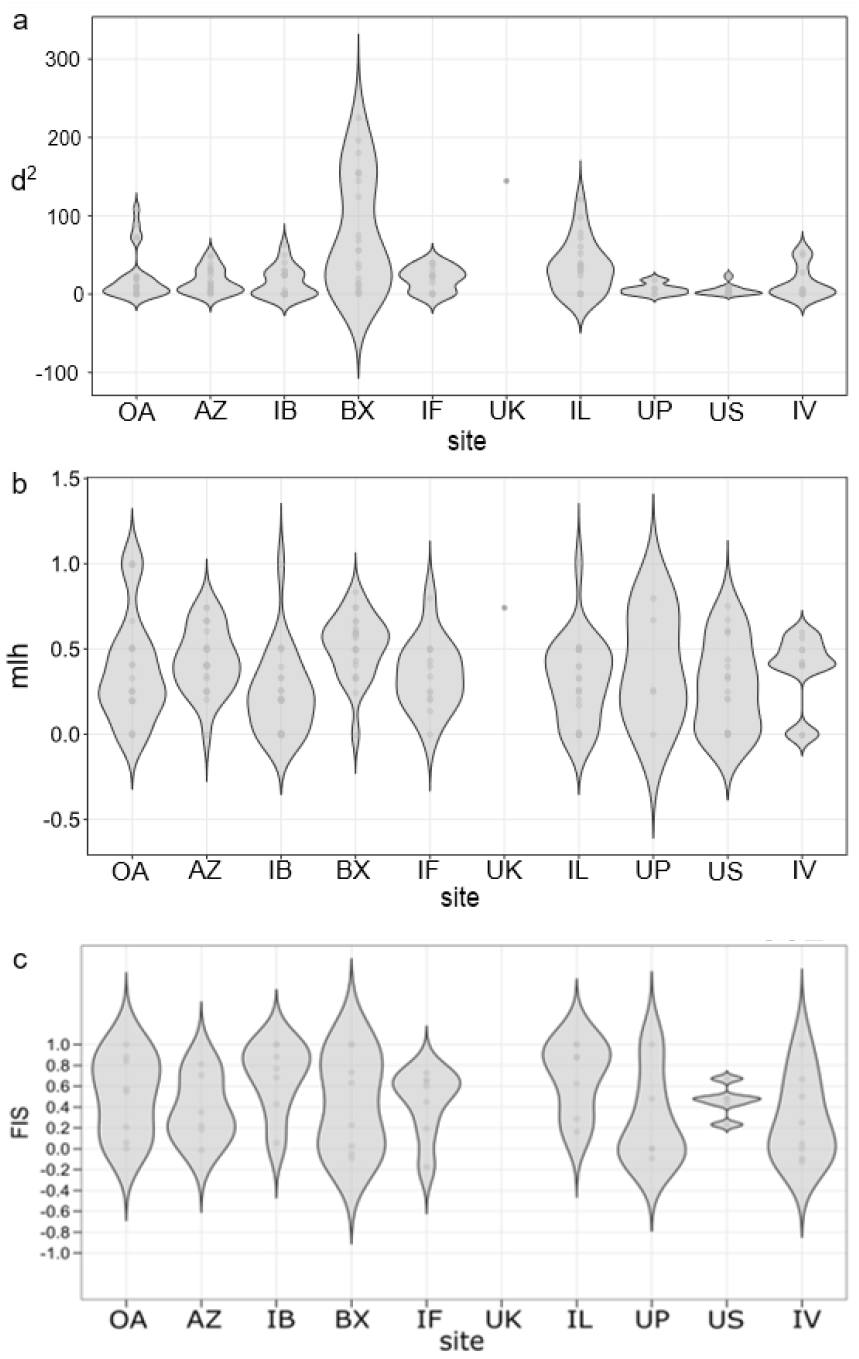
Violin plots with different genetic diversity indices for the ten sampling sites of 161 *Lissotriton vulgaris* individuals. Single points represent results of individuals. **a** Mean squared distance between alleles (d^2^); **b** Mean multi locus heterozygosity (MLH) and **c** Inbreeding coefficient (F_IS_). Abbreviations for different sites can be taken from Figure 1.

The PCA plot depicted a spatial separation of the sites, with the Innsbruck sites being distinct from the BX site. The Unterland and the Oberland sites were also close together, with some overlapping points (Figure 3, selected dataset Supplementary Figure S2).

**Figure 3.**
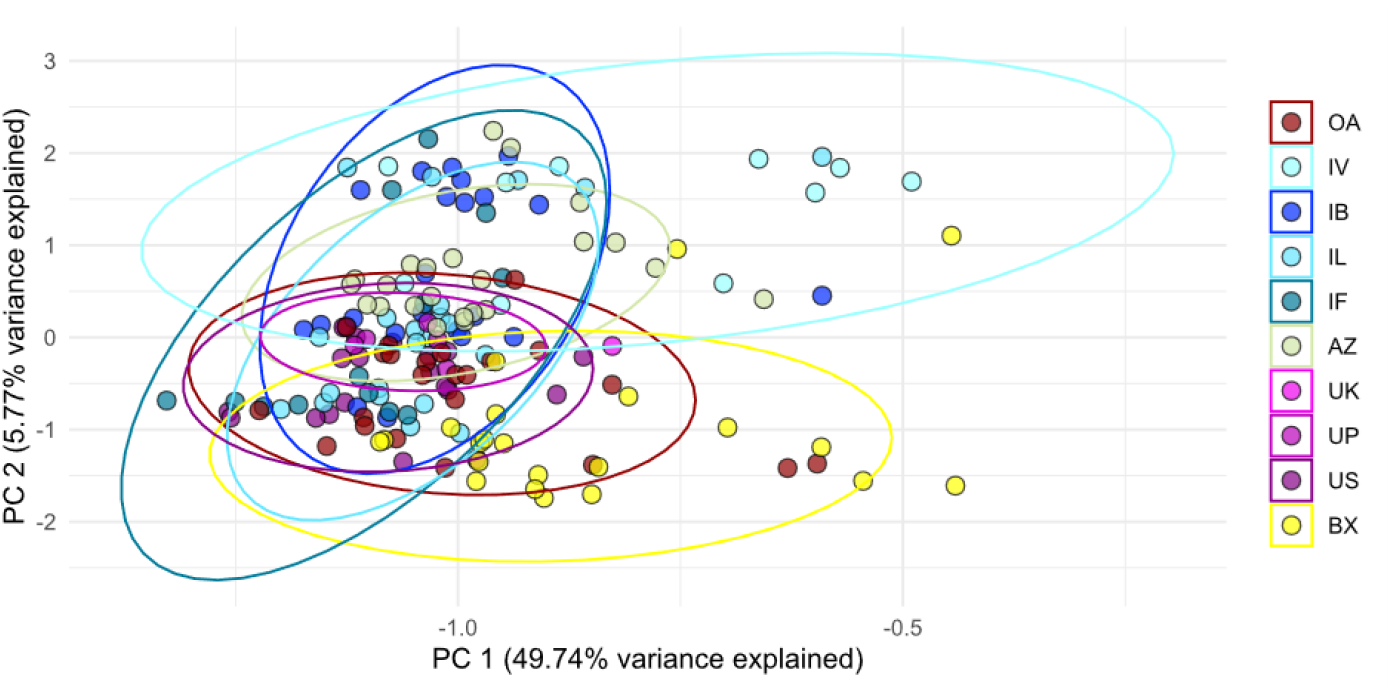
Principal component analysis (PCA) with 161 *Lissotriton vulgaris* individuals from ten different sampling sites. The first two axes explain 55.51% of the variance. The 95-percentile confidence interval is represented by an ellipse, with colors selected based on geographical location. The Innsbruck sites (blue) are predominantly located at the upper y-axis, where it looks like BX is more segregated at the lower edge. Abbreviations for the different sites can be taken from Figure 1.

The Bayesian clustering result depicted four populations. The population differentiation indicated three populations in North Tyrol and a different population for the subspecies *Lissotriton vulgaris meridionalis* in Brixen (Figure 4, Supplementary Figure S3-6).

**Figure 4.**
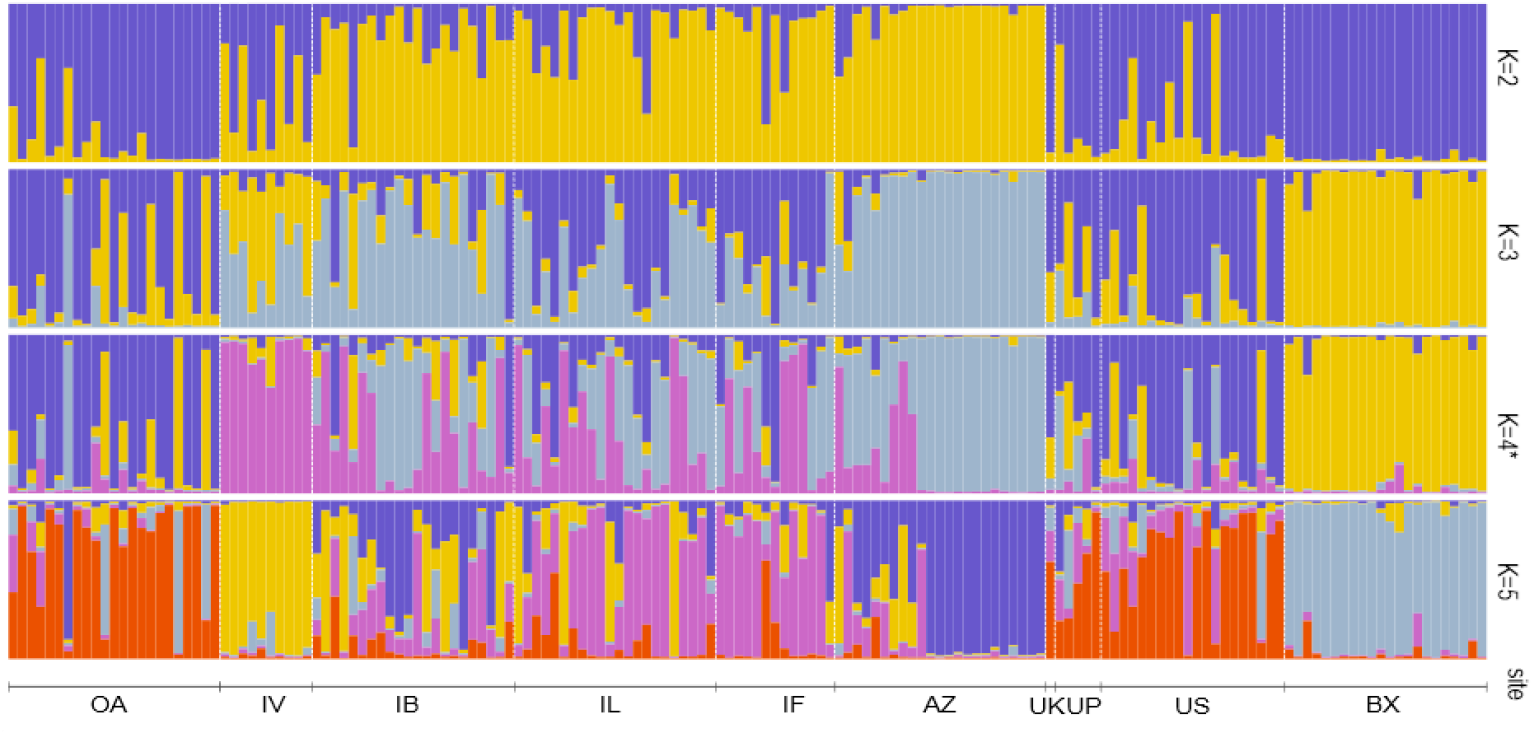
Concluded population structure of *Lissotriton vulgaris* for 161 individuals and ten sampling sites, generated using an admixture approach in STRUCTURE with 1,000,000 generations burn-in, and 2,000,000 MCMC generations and 10 replicates. K ranges from 2 to 5, with K=4* being the most likely population division. Two different clusters are shown for the AZ and IV sites, with mixture events occurring in the Innsbruck sites (IB, IL, IF). The other Tyrolean sites (OA, UK, UP, US) are categorized into the same cluster, indicating mixture events. BX with the subspecies *Lissotriton vulgaris meridionalis*, is clearly a distinct cluster. Abbreviations for different sites can be taken from Figure 1.

## Discussion

The objective of this study was to gain a better understanding of the current population genetic status of *L. vulgaris* in the Tyrolean Inn Valley by using non-invasive skin swabs instead of toe or tail clippings. Additionally, we aimed to better understand the potential influence of allochthonous individuals on autochthonous populations. We examined population genetics using microsatellite markers to clarify the genetic dynamics of native populations in comparison with artificially introduced populations. We further tested a captive population in a zoological garden and sampled individuals of the subspecies *L. vulgaris meridionalis* in Brixen, Italy, to calibrate our results. With no previous microsatellite study done on this species in Austria, we aimed at new insights into the local newt populations.

### Challenges and Considerations in Non-Invasive Genetic Sampling

The methodology we used, non-invasive skin swabs, presented notable challenges in acquiring genetic data. Of the eleven established microsatellite markers (Buono et al. 2018), three (Lm_528, Lm_749, LVG-210) were excluded due to unsuccessful gene scoring results (mismatching alleles in PCR replicates or monomorphism). We suspect that the skin swab technique was responsible for the high error rate across all microsatellite markers and individuals. We faced challenges similar to those described by Prunier et al. (2012), Müller, Lenhardt, and Theissinger (2013), such as inconsistencies in genotyping and allelic mismatch. In the study by Prunier et al. (2012), many failures were attributed to the dry skin of the frogs, especially during mating season, which additionally may have caused cross contamination. In our study of newts, we could exclude both excessively dry skin and significant cross-contamination, since the newts were caught individually out of the water during breeding season. Our decision to use ventral skin swabbing, instead of dorsal, could have negatively impacted our DNA yield (Prunier et al. 2012). Considering other potential sources of error in skin swabs, such as the influence of skin microbiota or specific alkaloids (Amézquita et al. 2017; Assis et al. 2017; Ringler 2018), it is likely that buccal swabs remain the more promising non-invasive method for extracting DNA from newts (Broquet et al. 2007). However, skin swabs cause potentially less stress and harm to the animal (Müller et al. 2013). From our eight remaining markers, loci Tv12 and Lm_013 had only five and six alleles, respectively (Table S1). Due to difficulties with the microsatellite markers and unequal individual numbers across sites, interpretation must be done with caution. Nevertheless, population genetic predictions can be made, as previously suggested by Buono et al. (2018).

### Genetic Diversity and Population Structure of Lissotriton vulgaris in the Tyrolean Inn Valley

The analysis of 161 *L. vulgaris* samples across ten sites revealed a diverse demographic composition with 87 females, 54 males, and 20 juveniles. Because of genotyping issues, analyses were done with the complete and a selected dataset (including just individuals with at least four genotyped loci). The comparison of these two datasets, showed overall consistent populations trends, indicating that the observed patterns are reliable despite the higher proportion of missing data in the full dataset (selected dataset Table S2-3, Figure S1-2,5-6).

We observed large variations in genetic diversity across the sites sampled, with mean observed heterozygosity (Ho) values ranging from 0.21 to 0.52 (Table 1). Ho values vary between the complete and selected dataset, particularly for individuals from AZ. Extreme values in some cases (e.g., in AZ) distorted mean values and can bias interpretation, which could be explained by including individuals with at least one locus present in the complete dataset. This was also observed for MLH (Figure 2). Therefore, the median observed heterozygosity (Median Ho), should be used for interpreting the data. In the selected dataset, especially the median Ho values, puts the Ho results into perspective, and the overall parameters align better at this site. This identifies BX as the site with the highest median Ho values in both the complete and selected datasets (Table 1, Table S2). Pairwise F_ST_ values ranged from 0.04 between IF and IL to 0.31 between OA and AZ (Table 2). AZ showed the highest differentiation from all other sites with higher values compared with our outgroup BX. This differentiation was also evident in the Bayesian cluster analysis, in which AZ stands out as a separate group of four in total (Figure 4). Since there is, to our knowledge, no population genetic study on *L. vulgaris* specifically in Austria, a careful comparison is only possible with other analysis done on this topic (Buono et al. 2018; Lark Davis 2021). These studies on Italian and Swedish newt populations experienced similar ranges of genetic diversity indices, with our Ho values having smaller extremes. The differences we observed among local populations are expected for *L. vulgaris*, as summarized in detail by Kalezic and Tucic (1984). It was concluded that there are substantial genetic differences between the populations due to limited gene flow, natural selection, and genetic drift, which are occurring independently at different sites. Moreover, *L. vulgaris* is philopatric, with its main dispersal happening through juvenile migration, and displays homing behaviour (Gill 1978; Fusillo et al. 2021). All of this, combined with often small effective population sizes, can lead to substantial genetic differences among newt populations. The Tyrolean Inn Valley, with its fragmented aquatic habitats, likely exacerbates these natural tendencies, shaping the distinct population structure we observed.

From our results, we conclude that the Unterland populations are in a more stable population genetic state than the Innsbruck sites. Both UP and US serve as our autochthonous references and are under federal protection, having been surveyed previously (Landmann and Fischler 2000; Schmidtler and Schmidtler 2001; Pagitz and Huemer 2018). These sites exhibit higher allelic numbers and higher median Ho values compared with other North Tyrolean sites (Table 1). The Bayesian cluster analysis groups all Unterland locations with OA, despite the low likelihood of genetic exchange between the sites. The clustering results align with relatively low F_ST_ values (Figure 4 and Table 2). However, caution is needed in interpreting pairwise F_ST_ values between the Unterland sites themselves in relation to OA, due to negative lower 95% confidence intervals (Meirmans and Hedrick 2011; Lee 2016).

### Anthropogenic Influences on Population Genetic Status

Our findings suggest that human activity is shaping the genetic landscape of *L. vulgaris* in the study area, especially among introduced populations. The Oberland site (OA), established in 2010 with the construction of the Area47 leisure park, represents a relatively young population. Its allelic richness (2.9), total alleles (Na=39), and median Ho (0.23) suggested a stable population (Table 1). Bayesian clustering arranged OA individuals with the Unterland sites (UK, UP, US). Given the lack of documented local autochthonous populations nearby (Landmann and Fischler 2000), we infer that OA was likely colonized by individuals introduced via aquatic plants or other materials brought into the artificial water body, underscoring the role of unintentional human-mediated dispersal.

Further, a genetic exchange between the Innsbruck sites, with a clear population difference between individuals from AZ and IV could be seen. The very low number of alleles (Table 1) and low d^2^ values (Figure 2) in AZ individuals, are likely reinforced by prolonged genetic isolation within the zoo. The results in Fig. 2 demonstrated the challenge posed by the data, as some markers had clustered values of zero or one, which again could bias the interpretation (as described in the paragraph before). However, the patterns seen suggest a tendency for inbreeding for multiple sites. In the AZ population, pronounced population differentiation is evident, supported by increased pairwise F_ST_ values between AZ and all other sites (Table 2) and a visible distinction in the Bayesian cluster analysis (Figure 4), despite the formal origin of AZ individuals from the autochthonous population of Kramsacher Loar (UK), retrieved in 1987 (Glaser 2008). Genetic mixture events can be observed in individuals from AZ, with the other Innsbruck sites IB, IL and IF (Figure 4) and seem to reflect the introduction of AZ individuals to IB. This suggests that genetic specification within the AZ population may have occurred prior to these releases and highlights the importance of maintaining the genetic health of isolated amphibian species. While ex-situ conservation programs can be a an additional tool for conservation efforts, in-situ habitat protection and restoration should be the primary goal in addressing global amphibian extinction. (IUCN 2005; Zippel et al. 2011).

While AZ and IV themselves presented homogeneous clusters in the Bayesian cluster analysis, IB, IL, and IF were more similar to each other, suggesting an ongoing geneflow among the sampling sites with inputs from AZ and IV (Figure 4). Although IL, IB, and IF exhibited similar patterns in Bayesian cluster analysis, direct migration between IF and the other sites is unlikely due to significant geographical distance. These similarities may be attributed to the introduction of a small number of AZ individuals in the late 1980s to IF (Landmann and Fischler 2000). The genetic similarities observed in the Innsbruck sites may be due to the existence of small autochthonous populations in the Inn Valley that were previously overlooked, as well as to individuals intentionally and/or unintentionally introduced for example through eggs on water plants (Landmann and Fischler 2000).

Population differentiation can be observed at the IV site. These individuals most likely descend from a single founder pair of *L. vulgaris* from East Tyrol, introduced for conservation purposes (Glaser 2008). Although the IV site showed higher allelic numbers (Na) and median observed heterozygosity (Median Ho) compared with nearby sites, its distinct genetic profile was evident in the Bayesian clustering (Figure 4) highlighting the genetic consequences of such translocations. Our study emphasizes the need for caution when transferring non-local populations of the same species. This intervention to reduce the population decline by restocking with allochthonous individuals is questionable (Reid et al. 2025), since the influence of geographically allochthonous populations on salamanders has not been extensively researched.

### Subspecies Differentiation and Biogeographic Context

Since STRUCTURE analysis is sensitive to missing values in datasets (Pritchard et al. 2000; Pritchard and Wen 2002), we evaluated our results to ensure the accuracy by comparing the North Tyrolean newts with the subspecies *L. vulgaris meridionalis* in BX, Italy. Our study identified three populations in the Tyrolean Inn Valley and correctly identified one separated population of *Lissotriton vulgaris meridionalis* in BX, Italy (Fusillo et al. 2021). This distinct separation corroborates the accuracy of our analysis.

The BX population is genetically robust, characterized by the highest number of alleles (Na=55) among all sampled sites, an allelic richness of 3.1, and median Ho of 0.48 (Table 1). This subspecies, geographically separated from the North Tyrolean newts by the main Alpine ridge, was clearly distinguishable through high pairwise F_ST_ values (Table 2), a pronounced separation in the Principal Components Analysis (PCA; Figure 3), and distinct clustering in the Bayesian cluster analysis (Figure 4).

However, we found the highest F_ST_ values not between BX and all other sites, but between AZ and all sites. One possible reason is the sensitivity of F_ST_ statistics to highly divergent populations when using microsatellite markers, resulting in an underestimation of differentiation (Balloux and Lugon-Moulin 2002). Despite this, the consistent evidence from multiple genetic analyses (STRUCTURE, PCA, and high F_ST_ relative to northern populations) strongly supports genetic divergence between *L. vulgaris meridionalis* and the *L. vulgaris* populations investigated in the Inn Valley, underscoring the influence of significant historical biogeographic barriers.

### Conservation Implications and Management Strategies

Our study provides valuable data for the conservation management of *L. vulgaris* in the Tyrolean region and beyond. The Unterland populations (UP, US) appeared to be in a generally more stable population genetic state. Despite the relatively high values of the genetic diversity indices we used and the inferred gene flow, persistently low catch rates at UP and especially UK continue to raise serious concerns regarding the overall population level of *L. vulgaris* in Tyrol. Comparative studies including the known autochthonous population in Tyrol (UK) with newly reintroduced sites would offer meaningful insights into newt population dynamics in the region.

The genetic depletion observed in the zoological garden population (AZ) highlights the significance of maintaining genetic diversity in isolated amphibian species. While ex-situ conservation programs can support conservation efforts, our findings strongly emphasize that in-situ habitat protection and restoration must remain the central and most effective strategies for addressing the ongoing global amphibian extinction (IUCN 2005; Zippel et al. 2011). Furthermore, the genetic differentiation seen at the IV and AZ site reinforces a crucial conservation message: Caution is necessary when considering the transfer or introduction of individuals from non-local populations. Such actions carry risks beyond outbreeding depression, loss of local adaptation, and introduction of maladaptive traits, all of which can undermine long-term population viability (Reid et al. 2025). As highlighted by Warne and Chaber (2023), translocation projects frequently lead to “translocation significant disease incursions” (TSDIs), causing negative population growth or failure to establish. These risks stem from factors such as the origin of the animals (wild-caught or captive-bred), the type of pathogen, and the host species. Our study exemplifies the necessity of comprehensive, standardized disease and genetic risk assessments as a mandatory prerequisite for any wildlife translocation project.

## Conclusion

This study provides new insights into the population genetics of North Tyrolean *L. vulgaris*. Three distinct population clusters were identified in North Tyrol, and differences between allochthonous and autochthonous populations were found. We found evidence of genetic depletion in captive individuals residing in a zoological garden. This highlights the importance of considering genetics in breeding programs before introducing them into the wild. The observed variations and population differentiation among sampling sites emphasize the significance of protecting as many populations as possible. The DNA extraction method of skin swabs did not yield the expected results when used for microsatellite genotyping. Further methodological studies on microsatellite markers in combination with skin swabbing would be beneficial for detailed population genetic studies of *L. vulgaris*. This study on Tyrolean *L. vulgaris* populations may serve as a starting point for the population genetic analysis of newts in Austria.

## Acknowledgements

Thanks to Bastian Angerer, Armin Leitner, Iris and Julia Schlick-Steiner, Miguel Steiner, Elisabeth and Marlies Stonig for field assistance, and to Florian Reischer and Elisabeth Zangerl for support in the laboratory. We are also grateful to the Alpenzoo Innsbruck and curator Gernot Pechlaner for the opportunity to sample captured individuals and to the authorities in Tyrol and South Tyrol for approval of our sampling.

