## SupplementaryMaterial for "Population structure and genetic diversity of smooth newts (*Lissotriton vulgaris*) in North Tyrol, Austria: influences of allochthonous individuals and conservation implications"

**Supplementary material**

**Tables**

**Table *S*1**: Eight microsatellite *markers* used in the analysis of 161 Lissotriton vulgaris individuals. The different alleles found are listed below the corresponding marker (allele) along with the frequency of occurrence (n). Marker Tv12 has the fewest alleles, with one dominant allele (237), *followed by* marker Lm_013 with six different alleles. Marker Lm_749 has the highest number of different alleles (22).

| **Th09 allele; n** | **Tv12 allele; n** | **Tv3Ca9 allele; n** | **Lm_749 allele; n** | **Lm_870 allele; n** | **Lv_542 allele; n** | **Lm_013 allele; n** | **Tv3Ca19 allele; n** |
|---|---|---|---|---|---|---|---|
| 185; 26 | 221; 7 | 116; 5 | 200; 8 | 144; 9 | 167; 20 | 166; 4 | 127; 5 |
| 189; 11 | 231; 2 | 120; 5 | 202; 8 | 156; 1 | 169; 5 | 168; 164 | 129; 1 |
| 201; 2 | 231; 1 | 122; 69 | 206; 4 | 160; 2 | 171; 18 | 170; 10 | 131; 28 |
| 205; 15 | 237; 109 | 124; 22 | 210; 4 | 168; 1 | 173; 6 | 178; 8 | 139; 4 |
| 209; 7 | 239; 61 | 126; 1 | 214; 20 | 172; 1 | 175; 13 | 180; 7 | 141; 43 |
| 213; 11 | | 130; 8 | 218; 3 | 176; 9 | 177; 14 | 216; 3 | 151; 25 |
| 217; 4 | | 138; 37 | 222; 43 | 180; 2 | 185; 4 | | 153; 3 |
| 221; 6 | | 142; 5 | 226; 18 | 184; 10 | 187; 4 | | 155; 2 |
| 225; 9 | | 144; 120 | 230; 23 | 188; 1 | 191; 1 | | |
| 229; 32 | | 146; 10 | 234; 29 | 192; 24 | 197; 1 | | |
| 233; 24 | | 148; 1 | 238; 11 | 196; 30 | 199; 2 | | |
| 237; 10 | | 150; 2 | 242; 22 | 200; 2 | | | |
| 241; 2 | | 154; 1 | 246; 4 | 204; 23 | | | |
| 245; 9 | | | 250; 1 | 208; 5 | | | |
| 247; 1 | | | 254; 3 | 212; 8 | | | |
| 251; 9 | | | 258; 1 | 216; 2 | | | |
| | | | 262; 7 | 220; 4 | | | |
| | | | 266; 1 | 224; 2 | | | |
| | | | 282; 6 | | | | |
| | | | 310; 9 | | | | |
| | | | 314; 3 | | | | |
| | | | 322; 2 | | | | |

**Table *S*2:** Selected dataset with 121 individuals: Sampling sites (site), number of sampled Lissotriton vulgaris individuals (N), allele number (Na), mean expected heterozygosity (He) , mean observed heterozygosity (Ho), 95% confidence interval (Ho Median (95%CI)), number of microsatellite marker used for calculation of He and Ho (n) and Bartlett's K square test of variance, as a proxy of differences between He and Ho (Bartlett's $K^2$) with p values in brackets. Calculations for UK *were* not possible. AZ results for He and Ho should not be considered, since Bartlett's $K^2$ is significant. Abbreviations for different sites can be taken from Figure 1.

| Site | N | Na | He (± SD) | Ho (± SD) | Ho Median (95%CI) | n | Bartlett's $K^2$ |
|---|---|---|---|---|---|---|---|
| OA | 18 | 35 | 0.58 (±0.22) | 0.35 (±0.33) | 0.24 (0.13-0.53) | 8 | 1.04 (0.31) |
| IV | 9 | 27 | 0.59 (±0.17) | 0.48 (±0.30) | 0.44 (0.33-0.60) | 8 | 2.04 (0.15) |
| IB | 12 | 24 | 0.61 (±0.25) | 0.21 (±0.24) | 0.13 (0.05-0.33) | 7 | 0.01 (0.92) |
| IL | 15 | 23 | 0.61 (±0.25) | 0.23 (±0.25) | 0.13 (0.05-0.39) | 7 | 0.00 (0.99) |
| IF | 12 | 28 | 0.53 (±0.23) | 0.33 (±0.22) | 0.29 (0.21-0.42) | 8 | 0.01 (0.92) |
| AZ | 17 | 19 | 0.64 (±0.26) | 0.35 (±0.21) | 0.34 (0.21-0.46) | 6 | 0.25 (0.62) |
| UK | 1 | 7 | | | | | |
| UP | 3 | 16 | 0.61 (±0.34) | 0.44 (±0.46) | 0.33 (0.08-0.83) | 6 | 0.43 (0.51) |
| US | 16 | 36 | 0.59 (±0.39) | 0.31 (±0.26) | 0.31 (0.12-0.44) | 7 | 0.87 (0.35) |
| BX | 18 | 53 | 0.68 (±0.23) | 0.48 (±0.4) | 0.51 (0.15-0.81) | 8 | 1.97 (0.16) |

**Table *S*3**: Pairwise fixation index ($F_{ST}$) values between ten different sampling sites of 121 Lissotriton vulgaris individuals and eight microsatellite markers. Small numbers below the $F_{ST}$ values represent the lower and upper 95% confidence interval. Values that exceed 0.05 suggest low genetic connectivity between the sit*e*s. AZ shows the highest genetic isolation with all sampling sites. Abbreviations for different sites can be taken from Figure 1.

| | OA | AZ | IB | BX | IF | UK | IL | UP | US |
|---|---|---|---|---|---|---|---|---|---|
| **AZ** | **0.28**<br>0.10; 0.44 | | | | | | | | |
| **IB** | 0.14<br>-0.01; 0.29 | **0.12**<br>0.03; 0.21 | | | | | | | |
| **BX** | **0.12**<br>0.06; 0.17 | **0.28**<br>0.14; 0.42 | **0.17**<br>0.08; 0.28 | | | | | | |
| **IF** | **0.11**<br>0.02; 0.19 | **0.20**<br>0.09; 0.32 | 0.06<br>-0.07; 0.17 | **0.18**<br>0.12; 0.23 | | | | | |
| **UK** | 0.07<br>-0.04; 0.19 | **0.14**<br>0.01; 0.30 | 0.03<br>-0.01; 0.08 | 0.04<br>-0.09; 0.14 | **0.10**<br>0.02; 0.22 | | | | |
| **IL** | **0.10**<br>0.04; 0.15 | **0.17**<br>0.07; 0.25 | 0.05<br>-0.03; 0.12 | **0.14**<br>0.08; 0.18 | **0.05**<br>0.00; 0.10 | 0.07<br>-0.01; 0.16 | | | |
| **UP** | 0.04<br>-0.19; 0.06 | **0.16**<br>0.03; 0.35 | 0.05<br>-0.19; 0.06 | 0.01<br>-0.14; 0.1 | 0.01<br>-0.16; 0.13 | 0.08<br>-0.32; 0.06 | 0.05<br>-0.06; 0.11 | | |
| **US** | 0.02<br>-0.05; 0.07 | **0.25**<br>0.05; 0.47 | 0.14<br>-0.04; 0.38 | **0.11**<br>0.04; 0.19 | 0.06<br>-0.03; 0.14 | 0.06<br>-0.08; 0.24 | **0.07**<br>0.03; 0.12 | 0.05<br>-0.18; 0.04 | |
| **IV** | **0.18**<br>0.04; 0.35 | **0.25**<br>0.11; 0.40 | 0.10<br>-0.01; 0.23 | 0.16<br>-0.00; 0.33 | **0.17**<br>0.08; 0.27 | 0.02<br>-0.17; 0.23 | **0.13**<br>0.06; 0.21 | 0.03<br>-0.16; 0.13 | **0.20**<br>0.03; 0.42 |

**Figures**

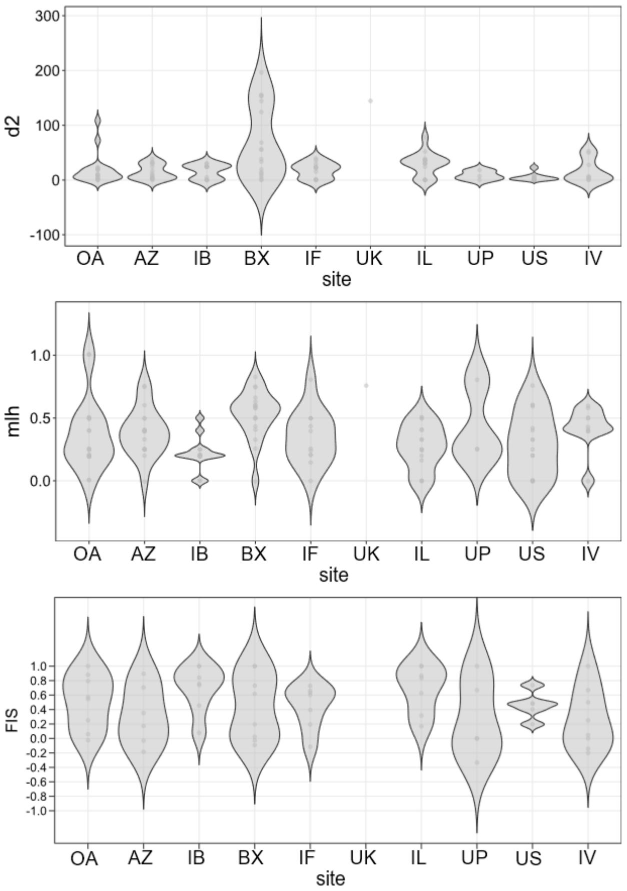


**Figure *S*1:** Violin plots with different genetic diversity indices for the ten sampling sites of 121 Lissotriton vulgaris individuals. Single points represent results of individuals. **a** Mean squared distance between alleles ($d^2$); **b** Mean multi locus heterozygosity (MLH) and **c** Inbreeding coefficient ($F_{IS}$). Abbreviations for different sites can be taken from Figure 1.

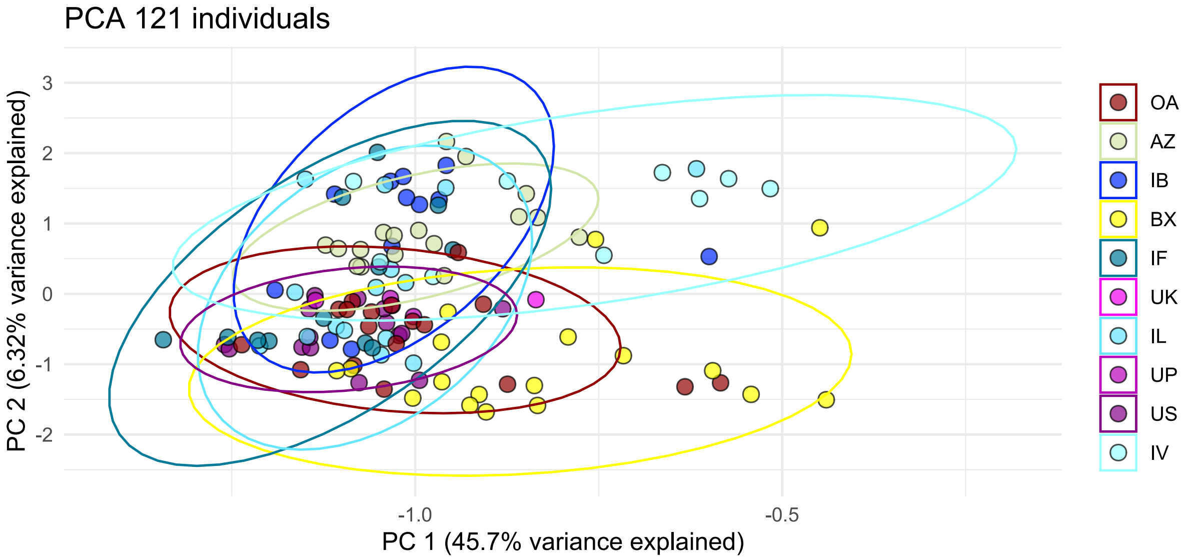


**Figure** ***S*****2**: Principal component analysis (PCA) with 121 Lissotriton vulgaris individuals showing ten different sampling locations. The 95-percentile confidence interval is represented by an ellipse, with colors selected based on geographical location. The Innsbruck sites (blue) are predominantly located at the upper y-axis, where it looks like BX is more segregated at the lower edge. Abbreviations for the different sites can be found in Figure 1.

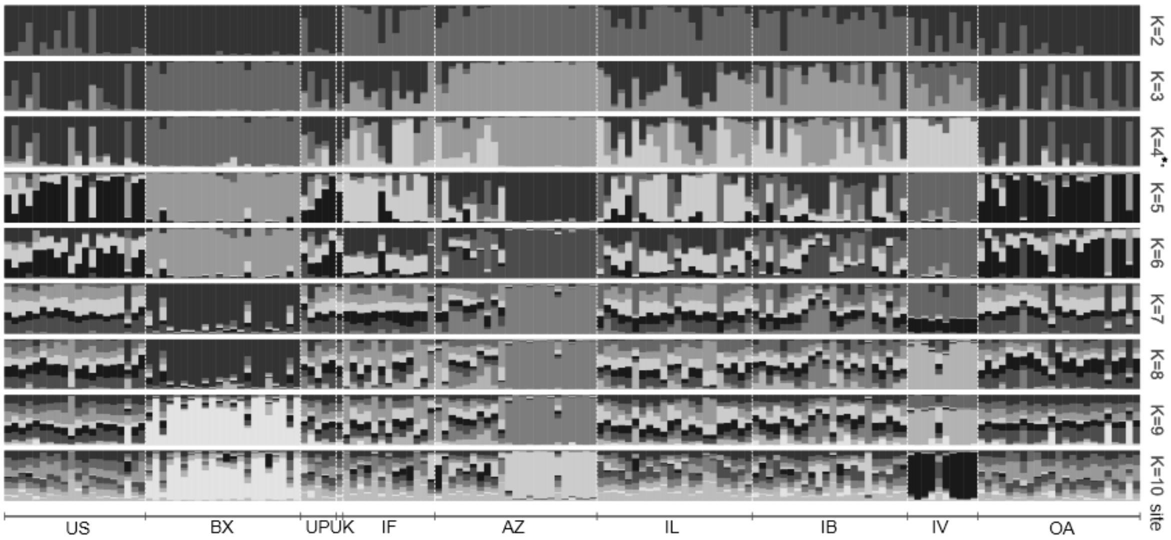


**S Figure 3**: Concluded population structure of Lissotriton vulgaris for 161 individuals and ten sampling sites, generated using an admixture approach in STRUCTURE with 1,000,000 generations burn-in, and 2,000,000 MCMC generations and 10 replicates. K ranges from 2 to 10, with K=4* being the most likely population division. Two different clusters are shown in AZ and IV sites, with mixture events occurring in the Innsbruck sites (IB, IL, IF). The other Tyrolean sites (OA, UK, UP, US) are categorized into the same cluster, indicating mixture events. BX with the subspecies Lissotriton vulgaris meridionalis, is clearly a distinct cluster. Abbreviations for different sites can be taken from Figure 1.

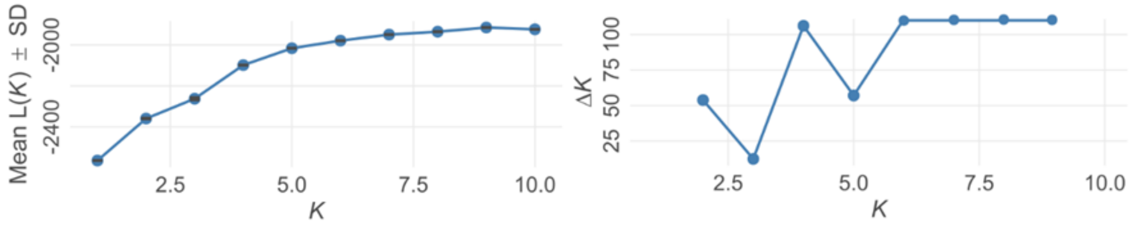


**Figure *S*4**: Suggested number of populations (delta K) for the complete dataset (161 individuals) with 1,000,000 generations burn-in, and 2,000,000 MCMC generations and 10 replicates for 161 Lissotriton vulgaris individuals and 8 loci. Delta K = 4 is the most likely number of population division.

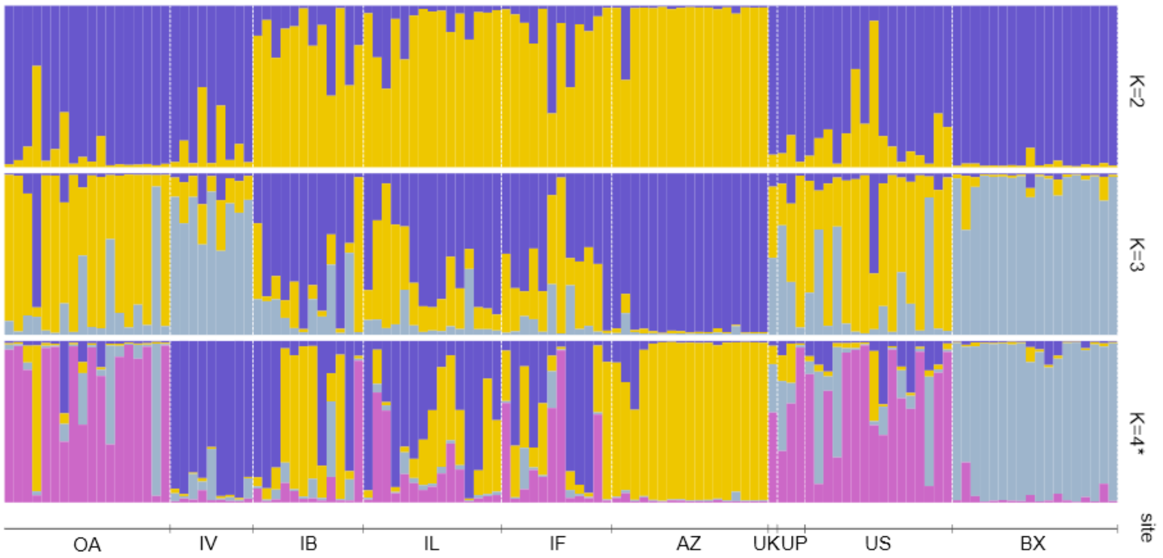


**Figure** ***S*****5**: Concluded population structure of Lissotriton vulgaris for 121 individuals and ten sampling sites. K ranges from 2 to 4, with K=4* being the most likely population division. Two different clusters are shown in AZ and IV sites, with mixture events occurring in the Innsbruck sites (IB, IL, IF). The other Tyrolean sites (OA, UK, UP, US) are categorized into the same cluster, indicating mixture events. BX with the subspecies Lissotriton vulgaris meridionalis, is clearly a distinct cluster. Abbreviations for different sites can be taken from Figure 1.

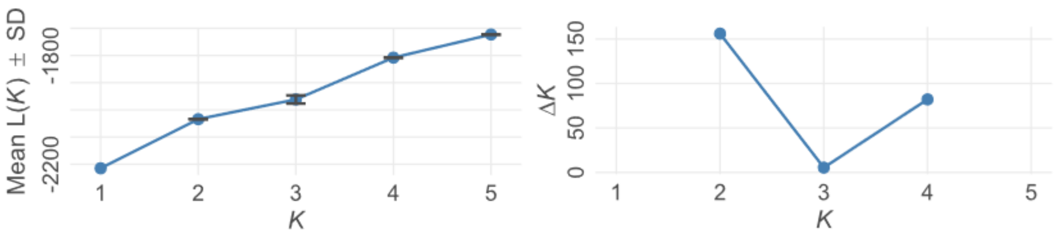


**Figure *S6***: Suggested number of populations (delta K) for selected dataset (121 individuals) with 1,000,000 generations burn-in, and 2,000,000 MCMC for 121 Lissotriton vulgaris individuals and 8 loci. Delta K = 4 is the most likely number of population division.
